# Molecular Basis of Urostyle Development: Genes and Gene Regulation Underlying an Evolutionary Novelty

**DOI:** 10.1101/2021.10.04.462674

**Authors:** Gayani Senevirathne, Neil H. Shubin

## Abstract

Evolutionary novelties entail the origin of morphologies that enable new functions. These features can arise through changes to gene function and regulation. One important novelty is the fused rod at the end of the vertebral column in anurans, the urostyle. This feature is composed of a coccyx and an ossifying hypochord, and both structures ossify during metamorphosis. We used Laser Capture Micro-dissection of these identified tissues and subjected them to RNA-seq and ATAC-seq analyses at three developmental stages in tadpoles of *Xenopus tropicalis*. These experiments reveal that the coccyx and hypochord have two different molecular signatures. ATAC-seq data reveals potential regulatory regions that are observed in proximity to candidate genes identified from RNA-seq. Neuronal (*TUBB3*) and muscle markers (*MYH3*) are upregulated in coccygeal tissues, whereas T-box genes (*TBXT*, *TBXT.2*), corticosteroid stress hormones (*CRCH.1*), and matrix metallopeptidases (*MMP1*, *MMP8*, *MMP13*) are upregulated in the hypochord. Even though an ossifying hypochord is only present in anurans, this ossification between the vertebral column and the notochord appears to resemble a congenital vertebral anomaly seen prenatally in humans, caused by an ectopic expression of the *TBXT*/*TBXT.2* gene. This work opens the way to functional studies that help us better elucidate anuran *bauplan* evolution.

## INTRODUCTION

Phenotypic and genotypic changes from an existing ancestral condition undergird the evolution of “key innovations” (Galis and Metz 2007). Phenotypic changes of a novel structure reflect changes in the corresponding genotypic/gene regulatory networks (Shubin, Tabin, and Carroll 2009; Tarazona et al. 2016; Tschopp and Tabin 2017; Wagner 2015). Previous studies have highlighted that the anuran (frog and toad) urostyle, composed of a coccyx and a hypochord, is morphologically unique from the rest of the vertebrates because of the contribution of an ossifying hypochord, and is therefore considered a structural novelty (Senevirathne et al. 2020; Handrigan and Wassersug 2007; Branham and List 1979; Kovalenko and Anisimova 1987; Kovalenko and Danilov 2006; Snell 2015). The coccyx, which is derived from the paraxial mesoderm, gives rise to the caudal vertebrae (Handrigan and Wassersug 2007; Sanchez and Sanchez 2013, 2015), which subsequently undergo endochondral ossification and fuse together during metamorphosis (Senevirathne et al. 2020). The amphibian hypochord, thought to be derived from either endoderm (Cleaver and Krieg 1998; Cleaver, Seufert, and Krieg 2000; Lofberg and Collazo 1997) or superficial mesoderm (Shook, Majer, and Keller 2004), is a thin embryonic rod, which degenerates in the rest of anamniotes during early embryonic development, but is retained only in frogs and undergoes endochondral ossification during the metamorphic climax (Handrigan and Wassersug 2007; Branham and List 1979; Kovalenko and Anisimova 1987; Kovalenko and Danilov 2006; Senevirathne et al. 2020; Snell 2015).

The ossifying hypochord, an apomorphic structure in anurans, occludes the dorsal aorta and is hypothesized to aid in rapid tail resorption (Senevirathne et al. 2020). We highlighted the phenotypic changes associated with the evolution of this structure in anurans and discussed how bones and cartilage, muscles, neurons form, and proposed how the hypochordal ossification has a role in the evolution of the anuran *bauplan* (Senevirathne et al. 2020). Despite being derived from two different populations of cells, both coccyx and hypochord undergo endochondral ossification during metamorphosis. Undifferentiated mesenchymal cells of the coccyx and embryonic hypochordal cells chondrify and ossify when the tadpole locomotion changes from an axial-driven mode to a limb-driven one. Ossification of the hypochord is rapid, usually ranging from 6-8 days. Apart from the cartilage and bone formation, the neuro-muscular skeleton is also remodeled. The muscles near the future caudo-pelvic region of the tadpole are remodeled during metamorphosis. The primary myotomes (Dorsalis trunci) remodel to form three different types of muscles (Longissimus dorsi, Coccygeoiliacus, Coccygeosacralis), where all three muscles attach to the coccyx. The axial motor neurons in the tail degenerate, and, at the same time, the spinal cord degenerates with the fusion of the coccyx and hypochord (Senevirathne et al. 2020).

Embryonic hypochord in anamniotes is known to have a function in remodeling the dorsal aorta, and the hypochord degenerates (except in anurans) after serving its purpose. Surprisingly, in mature tadpoles, CT scanning data revealed a possible role of the ossifying hypochord in remodeling the dorsal aorta as well. The posterior-most end of the hypochord appears to occlude the dorsal aorta, which could aid the rapid tail loss by cutting the blood supply to the tail (Senevirathne et al. 2020). Hence, we speculated that the ossifying hypochord has a role in the evolution of the anuran *bauplan*, and this could be a reason why it has been evolutionary favored in anurans for more than 200 million years (Shubin, Tabin, and Carroll 2009; Shubin and Jenkins 1995).

The phenotypic changes of the urostyle are well studied (Branham and List 1979; Kovalenko and Danilov 2006; Senevirathne et al. 2020); however, the molecular mechanisms underlying this unique structure have remained obscure to-date. Here, we investigate transcriptomic and gene regulatory networks in the developing urostyle by combining RNA-seq and ATAC-seq approaches. Using our previous morphology work (Senevirathne et al., 2020) as a framework to identify targeted cells, we used Laser Capture Microdissection to reveal the transcriptomics and epigenomics of the two tissue types, coccyx and hypochord.

Mesenchymal cells in vertebrates that undergo ossification have a conserved transcriptomic signature. Vertebrate ossification can be either endochondral or intramembranous, and a compendium of genes, transcription factors, intrinsic and external cues control ossification. During this process mesenchymal cells initially condense and commit to form osteoprogenitors (genes like *SOX2*, *RUNX2* are involved in this). Next, the osteoprogenitors differentiate to form preosteoblasts and osteoblasts (BMPs, FGFs, TGFß, and Wntß/catenin are involved in this (e.g., (Horowitz 2003; Karsenty 2008; Shen et al. 2014; Sodek and McKee 2000; Stein et al. 2003), and finally, mineralization and apoptosis of osteoblasts form mature osteocytes (e.g., (Horowitz 2003; Karsenty 2008; Shen et al. 2014; Sodek and McKee 2000; Stein et al. 2003).

Paraxial mesoderm-derived coccygeal cells are undifferentiated mesenchymal cells; they undergo chondrification and ossification prior to the initiation of the metamorphic climax (Handrigan and Wassersug 2007; Senevirathne et al. 2020) and could be following a similar gene regulatory network as connective tissues and bones in vertebrates. However, the ossifying hypochord initiates ossification at the onset of metamorphosis. The origin of amphibian hypochordal cells has been hypothesized to be from the endoderm (Cleaver and Krieg 1998; Cleaver, Seufert, and Krieg 2000; Lofberg and Collazo 1997), or the superficial mesoderm (Shook, Majer, and Keller 2004). Regardless of which germ layer it is derived from, hypochord undergoes endochondral ossification only in anurans. But the genes and gene regulatory regions that control the development of this structural enigma remain unknown. Here, we compare the gene expression patterns of coccygeal and hypochordal cells to identify similar/different pathways between the two tissue types, which are derived from two different cell populations.

Through this work, we address the following questions: Why does the hypochord only ossify in anurans? What are the similarities/differences between the hypochordal and coccygeal molecular pathways? Which genes switch on/off during metamorphosis? By identifying the underlying changes in the genes and gene regulatory networks, our work begins to shed light on the potential genotypic changes underlying a structural novelty.

## MATERIALS AND METHODS

Different stages of *Xenopus tropicalis* tadpoles were purchased from the National Xenopus Resource (NXR) at the Marine Biological Laboratory (MBL), Woods Hole, MA. Comparisons were made across three significant life-history stages to highlight the differences/similarities of genes and gene regulatory dynamics during metamorphosis. The developmental stages used for the experiments were as follows: before metamorphosis/prometamorphic stages (stage 56/57), at the beginning of the metamorphic climax (stage 60/61), and end of metamorphosis (stage 65/66). The tadpoles were euthanized using 0.2% aqueous tricaine methanesulfonate (MS-222), and the specimens were fixed in different fixatives or fresh tissues were taken according to each experiment. Tadpoles were staged according to Nieuwkoop and Faber (NF). The codes generated for the bioinformatics analyses are deposited in GitHub (https://github.com/GayaniSenevirathne/Senevirathne_et_al_RNAseq.git) and the raw sequences are available at NCBI (Accession numbers will be given upon acceptance).

### RNA-seq using spatial transcriptomics and Laser microdissection (LCM)

*Xenopus tropicalis* tadpoles at prometamorphosis (stage 56), beginning of the metamorphic climax (stage 60/61) and end of metamorphosis (stage 65/66) were selected as the targeted stages for the RNA-seq experiment (stages were selected based on the significant phenotypic changes that were seen at each stage during the urostyle development based on Senevirathne *et al*., 2020). All forceps, scissors, surgical blades and lab benches were cleaned with RNAse away and 100% ethanol prior to any RNA sequencing experiment. Tadpoles were euthanized using MS-222. The region where the urostyle forms (demarcated by the tenth and fourteenth myotomes; Senevirathne *et al*., 2020) was dissected under a Leica L2 light microscope on ice-cold 1x DEPC-treated PBS; all the dissections were done on ice to prevent RNA degradation. The dissected tissue was immediately transferred to ice-cold OCT and flash frozen in liquid nitrogen and stored at −80°C (for better RNA quality, the tissue blocks were processed the subsequent day). The frozen tissue blocks were sectioned using a Leica cryostat.

To carry out a transcriptomic survey during urostyle development, we adapted a spatial transcriptomic approach (using Laser capture microdissection). The two targeted tissue types, coccyx and hypochord, from three individuals at each developmental stage (prometamorphosis, beginning of metamorphic climax, and end of metamorphosis) were dissected from frozen sections (Fig. 1A). The myotomic boundaries were used as a way of identifying the targeted area to be dissected out. Cells of interest were identified based on Senevirathne *et al* (Senevirathne *et al*., (2020)). Prometamorphic (stage 56) sections of coccyx had undifferentiated mesenchymal cells around the spinal cord, and the hypochord had embryonic hypochordal cells ventral to the notochord. The RNA-sequencing protocol followed a spatial transcriptomics method (Geo-seq; (Chen et al. 2017)). Prior to sectioning, the cryostat, brushes, adjacent benches/tabletops, blades, and pencils/pens were cleaned using RNAse away and 100% ethanol. The tissue blocks were left inside the cryostat for 20 minutes, allowing them to equilibrate at −20° C (not doing this resulted in flaky sections or sections breaking when transferred onto the slides). The tissues were sectioned at 16 μM thickness on to PEN membrane 1.0 slides. Five–six sections were placed on each slide and were allowed to dry at room temperature for one minute before storing them at −80°C for further processing (samples that were <1 month old were used for sectioning; the yield of RNA was high when the slides were sectioned on the same day).

**Figure 1:**
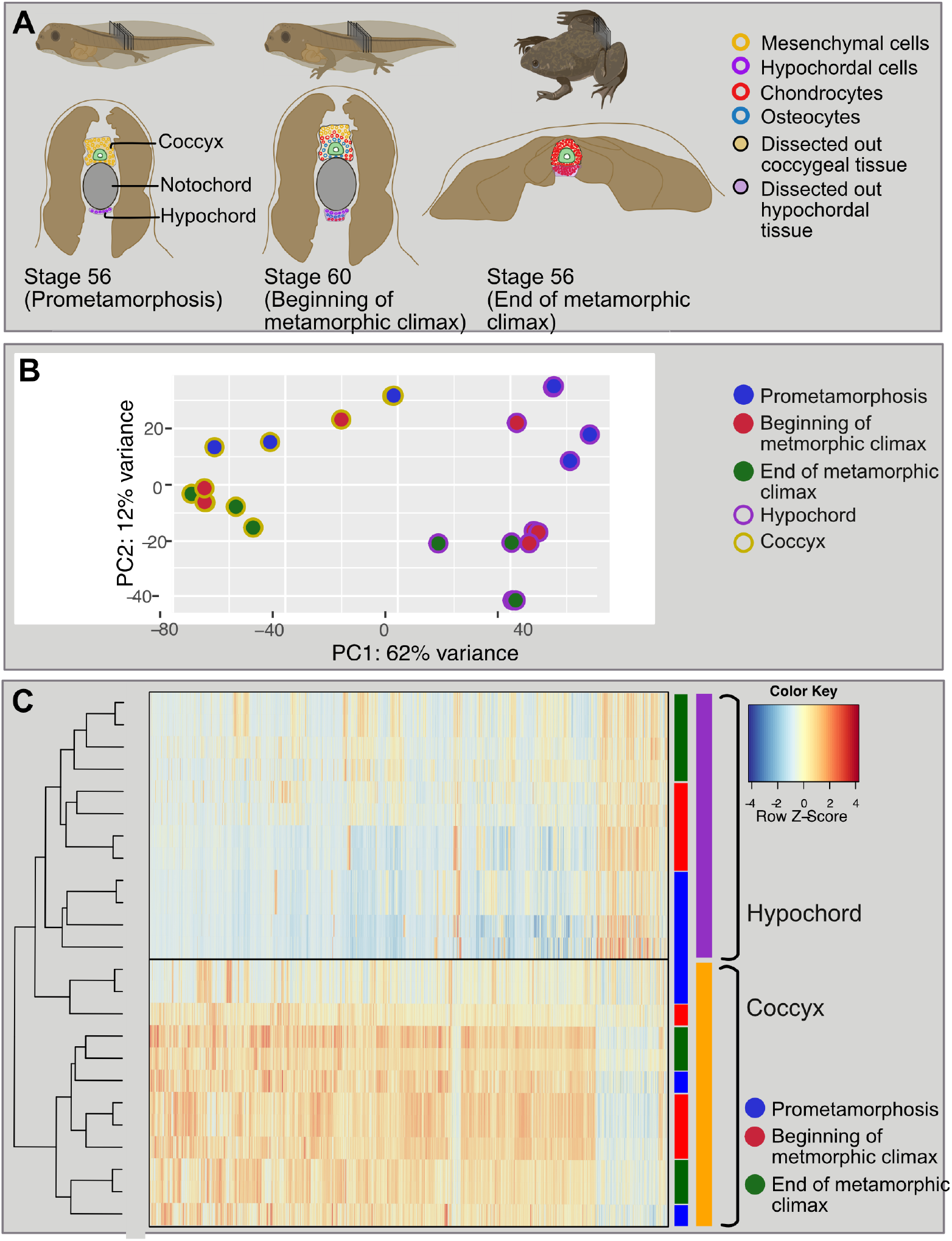
Changes in transcriptomics during anuran urostyle development. A. The experimental setup and the developmental stages used for Laser-capture microdissections. Ten sections of cryosections (16 mM each) were taken from three developmental stages (stage 56: prometamorphosis; stage 61: beginning of the metamorphic climax; stage 65: end of metamorphosis) and the coccygeal and hypochordal tissues were dissected out. B. Principal component analysis for the urostyle tissues (coccyx and hypochord) used for the transcriptomic assay (N=4 biological replicates per developmental stage). B. Principal component analysis of the log-normalized count data for all 24 samples. Each dot represents a tissue sample. C. Heatmap, highlighting the differentially expressed genes, compared between three developmental stages and two tissue types. The heatmap highlights that the two tissue types possess two distinct sets of genes.

On the day of the Laser capture microdissection, slides were removed from the freezer, thawed at room temperature for 2 minutes, and placed under an UV lamp for 2 minutes (UV helps the sections to adhere to the slide). Next, the slides were stained using Cresyl Violet to help visualize the cells. For this, slides were taken along an ethanol series, each wash was 30 seconds (100% ethanol, 70% ethanol, Cresyl Violet in 70% ethanol, and were dehydrated in 70%, 90%, and 100% ethanol). Slides were allowed to dry completely before moving to the next steps (this step was important to avoid humidity affecting the RNA quality (Ordway et al. 2009).

The dehydrated slides were processed via LCM with the following settings: aperture (10), speed (20) and energy (50). The hypochordal and coccygeal cells were identified (histological comparisons done in Senevirathne *et al*. (Senevirathne et al. 2020) were used as a reference) using the x10 eye piece and the dissections were done using the x20. Targeted cells were captured to an adhesive cap Eppendorf tube, with the cap consisting of 50 ul of the lysis buffer. Once the cells from coccyx and hypochord were collected (∼10,000 cells from 10 sections for each tissue type, 4 replicates were done for each stage, a total of 24 samples), 150ul of the lysis buffer was added to each tube and was left on ice for 20 minutes. RNA was extracted from the captured cells using the TAKARA NucleoSpin® RNA XS (Cat. No. 740902) kit with slight modifications (the filtration step was skipped). cDNA was generated using the SMART-Seq® v4 Ultra® Low Input RNA Kit for Sequencing (with the number of amplification cycles set to 18). cDNA was purified using Agencourt Ampure XP magnetic beads (Beckman Coulter) and were sequenced using the HiSeq PE100.

### Gene regulation and ATAC-seq

The same developmental stages that were used for the RNA-seq studies were taken, and the urostyle region was dissected out as a fresh chunk of tissue (morphological demarcations of the developing urostyle were decided based on Senevirathne et al. (Senevirathne et al. 2020)). The OMNI-ATAC-seq protocol was used to identify open chromatin regions in the developing urostyle (two replicates from each developmental stage, coinciding with the RNA-seq and morphological studies, were selected).

The tadpoles were anesthetized in MS-222, dissected on ice-cold 1X PBS and were mechanically crushed using a pestle (cleaned using 100% ethanol prior to this step) to obtain a homogenized sample (all these steps were done on ice to prevent degradation of proteins). Once a homogenized sample was obtained, cells were counted using the BioRad Tc20 automated cell counter. All samples consisted of 75,000–100,000 cells. The subsequent steps followed the OMNI-ATAC seq protocol (Buenrostro et al. 2015; Corces et al. 2017) with slight modifications using the Illumina Tagment DNA Enzyme and Buffer kit: cells were lysed in an ice-cold lysis buffer, followed by a transposition step using Tn5 Transposase, and DNA was purified using the Zymo DNA Clean and Concentrator. Purified DNA was amplified with 13 amplification cycles (the number of cycles were optimized by an additional qPCR step). Finally, the libraries were purified using the Zymo DNA Clean and Concentrator and were sequenced using the NovaSeq 2000 (100BP PE).

### RNA-seq analyses

Three stages were targeted for all the next-generation sequencing steps – Before metamorphosis (stage 56/57), beginning of metamorphosis (stage 60/61), and end of metamorphosis (stage 65/66). RNA from two different regions, coccyx and the hypochord, was extracted from three individuals for each stage (18 samples). 9 samples were run per lane, using paired end 100 bp reads on a llumina HiSeq 2000, at the Genomic core at the University of Chicago.

Sequence quality was checked using FastQC (version 0.11.9). (Please see Appendix A for sequence depth, Bioanalyzer results, and quality check files). *Xenopus tropicalis* reference genome v. 9.1 (Xenopus_tropicalis_v9.1.dna.toplevel.fa.gz) and transcript annotations were downloaded from Ensembl (www.Ensembl.org). Adapter sequences were trimmed using Cutadapt (version 1.8.1). Trimmed sequences were mapped using two approaches to compare the differentially expressed genes: 1. Normal alignment using HTSeq v.0.13.5 (Anders, Pyl, and Huber 2015) and Bowtie2 v.2.4.2 (Langmead and Salzberg 2012); 2. Pseudoalignment using Kallisto v.0.46.0 (Bray et al. 2016) were used to assess differentially expressed genes across tissues and developmental time points. Counts for HTSeq2 and Bowtie2 alignment files were obtained using HTSeq-counts computed for the *Xenopus tropicalis* v.9 annotations. Kallisto counts were also used as a comparison method. The subsequent steps are for the aligned transcripts obtained from the HT-seq2 step. The differential gene expression between the two tissue types (coccyx and hypochord), three developmental stages, and three biological replicates, were analyzed using the DESeq2 (Love, Huber, and Anders 2014) package (v.3.12) from Bioconductor. The dataset consisted of a total of 18 libraries (9 individuals, 3 replicates, 2 tissue types, 3 stages), differentially expressed genes were looked for either between stages (e.g., prometamorphosis *vs* beginning of metamorphic climax) or between the two tissue types (e.g., coccyx *vs* hypochord). A DESeq2 negative binomial generalized linear model was adapted, which has been highlighted in previous studies (Love, Huber, and Anders 2014) as a robust method for identifying differentially expressed genes (DEGs). DESeq2 package was used in R to normalize the reads, and the reads were subjected to variance stabilizing transformation using the “vst” function. A principal component analysis (PCA) was carried out using the DESeq2 function “plotPCA” to observe the clustering of the 18 samples. Hypochord and coccyx show considerable differences in cellular composition and differentiation (Senevirathne et al. 2020), and the gene expression profiles directly reflect this (Fig. 1C). A False Discovery Rate (FDR) value of <0.05 was used as the statistical significance threshold. DEG comparisons were depicted in three ways: prometamorphosis *vs* beginning of metamorphic climax; beginning of metamorphic climax *vs* end of metamorphosis; coccyx *vs* hypochord. The results of the DEG experiments were visualized in three main ways: 1. heatmaps were generated from the lists (Appendices C, D and Tables 3.1 and 3.2) of significant genes using the normalized values. Differences in expression data were visualized using z-scores calculated for each gene (=each row); 2. Volcano plots were drawn highlighting the up/down regulatory genes in the DEGs. Here, log-transformed p-values (y-axis) were plotted against the log2 fold change (x-axis); 3. Narrowed down gene symbols of the DEGs were used for GO enrichment analysis. The reactome web-based analysis tool was used to determine the overrepresentation of Reactome pathways where the up/down regulatory gene lists (Appendices C,D,E and F), genes within the intersections of the Venn Diagrams (drawn using the package “VennDiagram”) were given as inputs.

### ATAC-seq analyses

Adapter sequences were trimmed from the raw paired end 100-bp files using NGmerge (Gaspar 2018) and the trimmed sequences were aligned to the *X. tropicalis* reference genome v. 9.1 (Xenopus_tropicalis_v9.1.dna.toplevel.fa.gz) using Bowtie2. Duplicated reads were removed from the subsequent analyses using Picard (http://broadinstitute.github.io/picard/). Peaks were called using MACS2 (Zhang et al. 2008) (--nomodel --extsize 200 --shift -100 --nolambda) and Genrich (-e chrM -r -j). Two peak callers were used to compare the peaks, where Genrich’s “j” command specifically signifies the ATAC-seq mode. Irreproducible discovery rate (IDR) <0.01 was used as the threshold to screen the replicate samples. Here, the IDR method compares ranked peak lists to identify overlapping peaks. Finally, the peak files were directly uploaded to Integrative Genomics Viewer (IGV) and were visualized along with their respective. bam and bam index files.

### HCR *in-situ* hybridization

Targeted urostyle tissues were fixed in 4% PFA, dehydrated in a methanol series, and stored at −20° C until future use. On the day of sectioning, tissues were rehydrated using an ethanol series, rinsed in histosol, and subsequently, washed and mounted in paraffin. The microtome, brushes, bench/tabletops were cleaned using RNAse away and 100% ethanol and the tissue blocks were sectioned to obtain 12 uM-thickness paraffin sections. Paraffin sections can be stored at room temperature, indefinitely, until the day of staining.

For HCR *in-situ* hybridization (Yamaguchi et al. 2015) of paraffin sections (the protocol followed https://www.molecularinstruments.com/protocols with slight modifications), the sections were initially dewaxed using histosol, re-hydrated in ethanol, and treated with a Proteinase K/PBS solution to increase the tissues’ permeability. Prehybridization step was followed by the addition of the targeted probe (1 μM probe/100 ul of hybridization buffer) and leaving the slides in a 37° C incubator overnight. Next day, the slides were washed in the wash buffer and subjected to an amplification buffer with hairpins overnight. On the third day, the slides were washed using dilution a series of SSCT, mounted using Fluoromount G + DAPI, and visualized using a Zeiss LSM 710 confocal microscope. The results were analyzed using Fiji image analysis software.

## 3.4 RESULTS

### Disparity in gene expression profiles of the Coccyx and Hypochord

At the beginning of metamorphic climax (stage 60/61) both hypochordal and coccygeal cells underwent chondrogenesis and osteogenesis (dissected cells at this stage included immature chondrocytes, mature chondrocytes, osteocytes, mesenchymal cells, and extracellular matrix) (Fig. 1A and 2). At the end of metamorphosis, coccygeal and hypochordal cells completed ossification, and the majority of the cells consisted of osteocytes, osteoblasts, and mature chondrocytes. The two tissue types fuse at the end of metamorphosis, coinciding with the degeneration of the notochord. The total analysis consisted of 21458 genes, out of which 3286 genes exhibited considerable variation between the two tissue types across development (the FDR <0.05); both tissue types and the three timepoints were used as factors in the DESeq2 analysis where a binomial generalized linear model was implemented. Principal component analysis (PCA) revealed that the coccygeal and hypochordal samples generate two separate clusters (Fig. 3B), and a heatmap showed the two tissue types possess two different gene expression profiles (Fig. 3C). 3298 genes were differentially expressed between the urostyle and hypochord, whereas 1845, 385 and 3434 genes were differentially expressed between the prometamorphic *vs* beginning of metamorphic climax, beginning of metamorphic climax *vs* end of metamorphosis, and prometamorphosis *vs* end of metamorphosis, respectively. Among these DEGs, 2828 genes were significantly upregulated and 470 were downregulated in the coccygeal region compared to hypochord. During coccygeal development, several modifications happen around the areas of interest. The coccyx develops dorsal to the notochord and around the spinal cord, initially as two ossification centers, which later fuse together during metamorphosis. Concomitantly, muscles and neurons around the coccyx remodel. Primary myotomes remodel into secondary muscles and attach to the coccygeal bone. The spinal cord degenerates and axons project outwards from the coccygeal spinal foramina (Senevirathne et al. 2020). These phenotypic changes are reflected in the underlying gene regulatory networks. The majority of the upregulated genes in the coccygeal tissue samples are involved in differentiation and development of the nervous system (e.g., *NEUROD6*, *PRDM12*, *COCH*, *APBA2*) (Uittenbogaard, Baxter, and Chiaramello 2010; Rahman et al. 2020), or are genes that are expressed during skeletal muscle development (e.g., *ACTN2*) (Mills et al. 2001). Apart from these, the rest of the upregulated genes within coccygeal tissues are directly involved in chondrocyte/osteocyte differentiation (e.g., *RUNX2*, *COL9A1*, *SOX8*) (Fig. 3) (Youlten et al. 2020; Qin et al. 2020).

**Figure 2:**
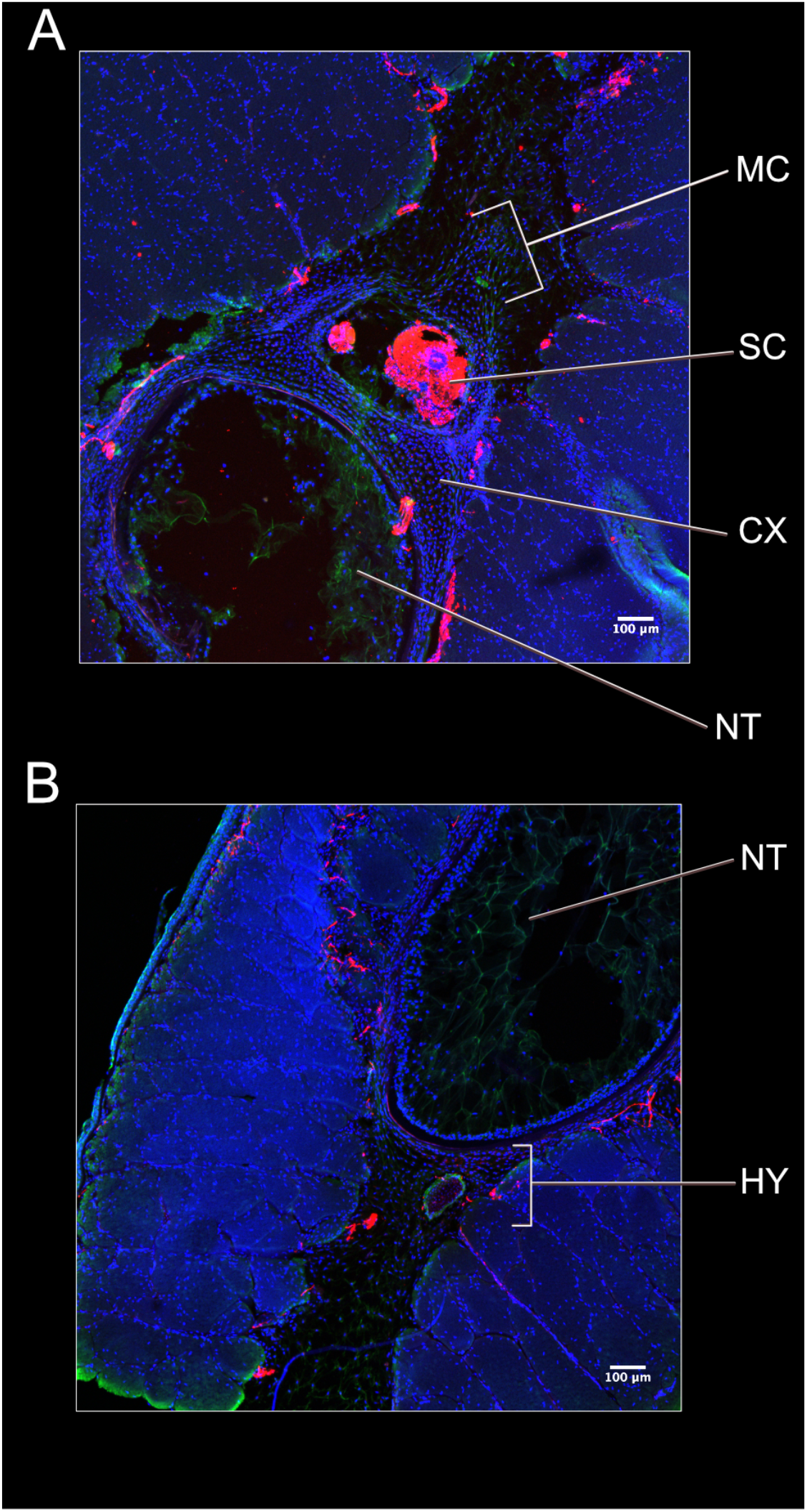
Comparison of the hypochordal and coccygeal sections before metamorphosis (stage 57). A. A transverse section across the coccyx, highlighting the aggregating mesenchymal cells around the spinal cord. B. A transverse section across the hypochord, highlighting the embryonic hypochordal cells ventral to the notochord and notochordal sheath. Nuclei stained in blue, using DAPI and neurons stained in red using acetylated tubulin.

**Figure 3:**
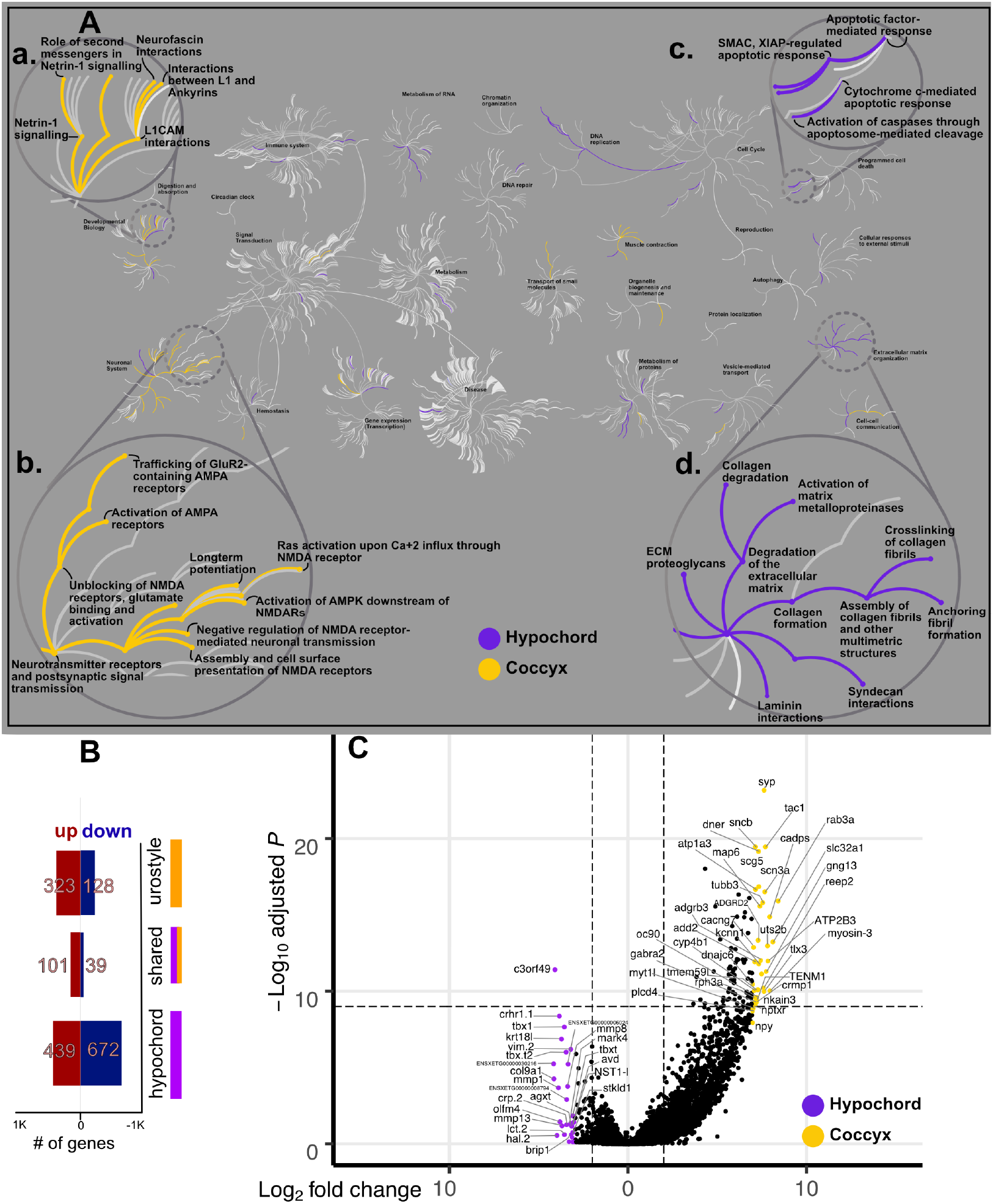
Comparative transcriptomic analysis of the two tissue types: coccyx and hypochord. A. A Reactome pathway analysis for up/down regulatory genes in coccyx vs hypochord; the central circles represent a top-level pathway, and the circles away from the center represents lower levels in each respective pathway. Zoomed-in sections of top-level pathways of Developmental Biology (Aa), Neuronal system (Ab), Programmed cell death (Ac), and Extracellular matrix organization (Ad) are shown. Overrepresented pathways (P < 0.05) are colored in yellow (coccyx) and purple (hypochord). Pathways that are not significant are shown in light gray lines. B. Most hypochordal genes are involved in organizing the extracellular matrix, whereas the majority of coccygeal genes are involved in neuronal remodeling and modifications. B. The total number of urostyle-responsive genes (FDR < 0.01) between hypochord and coccyx. C. Volcano plot showing differentially expressed genes across hypochord and coccyx during development (P < 0.05, FDR<0.01).

Embryonic hypochordal cells are thought to have an endodermal (Cleaver and Krieg 1998; Cleaver, Seufert, and Krieg 2000; Lofberg and Collazo 1997; Senevirathne et al. 2020) or a superficial mesodermal origin (Shook, Majer, and Keller 2004). Whether it is endoderm- or superficial mesoderm-derived, a cell population that is completely different from the sclerotomal cells (of the coccyx) forms the ossifying hypochord and contributes to the adult axial column. Hence, this unusual ossification of the hypochord, seen only in anurans (ranging from the myotome 10–14), is considered an apomorphic state, compared to the rest of the vertebrates. Embryonic hypochordal cells undergo chondrification and ossification as soon as the tadpole reaches its metamorphic climax. The transcriptomic assay between the two tissues revealed that hypochordal tissues express high concentrations of *TBX1*, *TBXT.1*, *TBXT.2*, and *HAND2* (Fig. 3C). T-box genes are involved in early mesodermal patterning and their expression has not been recorded in adult tissues before (explained in detail in a subsequent section; 3.4.3). Here, we hypothesize three possible scenarios: if the hypochordal cells are of endodermal origin, the increased *TBXT*/*TBXT.2* could be initiating a cell-fate switch from endoderm-to-mesoderm (there are some instances where T-box genes have been recorded to enable a cell-fate switch e.g., (Chapman et al. 2003)). To the best of our knowledge, there are no other studies looking into the possibility of an endoderm-derived tissue undergoing ossification. Secondly, if the hypochordal cells are superficial mesoderm derived, *TBXT* and *TBXT.2* could be activating the downstream targets involved in cellular matrix organization (seen by the up-regulated expression patterns of *MMP1*, *MMP8*; Fig. 3) and chondrification. Thirdly, another possibility is that the ossification of the hypochord could resemble an epithelial-to-mesenchymal transition (EMT). During EMT, the epithelial cells adapt a morphology similar to fibroblasts and acquire migratory properties (at the same time the epithelial cells lose adhesion to the surrounding extracellular matrix) (Radisky 2005). Several studies hypothesize how Brachyury (TBXT/TBXT.2) plays a pivotal role in EMT, where overexpression of Brachyury would induce mesenchymal properties, and reduce epithelial properties, in the migrating epithelial cells (Behr et al. 2005; Fernando et al. 2010). This phenomenon has led to abnormal ossifications in the vertebral column (i.e., vertebral column chordomas, where some are observed between the notochord and vertebral column) (Chen et al. 2020; Vujovic et al. 2006; Zhu, Kwan, and Mackem 2016). There are endothelial cells lying between the embryonic hypochord and endoderm (where the dorsal aorta runs between these two tissues) (Senevirathne et al. 2020). Hence, the increased expression of *TBXT*/*TBXT.2* in hypochordal cells could potentially lead to increased mesenchymal properties and eventually activate chondrifying and ossifying genes.

T-box genes have already been identified as being pivotal components in the differentiation of the posterior axial column (Chen et al. 2020; Cunliffe and Smith 1994, 1992; Gentsch et al. 2018; Ghebranious et al. 2008; Hayata et al. 1999; Hotta et al. 2000; Messenger et al. 2005; Schulte-Merker and Smith 1995; Vujovic et al. 2006; Wan et al. 2016), and seem to be playing a role in hypochordal ossification as well. However, all the three scenarios explained above, would require the activation of T-box genes at the onset of metamorphosis because of extrinsic/intrinsic signals, which could be either hormonal or environmental.

The pelvic region undergoes dramatic changes during metamorphosis, and this period is thought to represent the developmental stage that is most susceptible to predation. The underlying stress of the remodeling tissues and hormonal responses can also be seen by the increased expression of *CRCH.1* (corticosol steroid stress hormones), having a normal hormonal response to stress. Other than these genes, the hypochord also expresses significant concentrations of *VEGF* and *HAND2*. These two genes are involved in vascular development and can also be seen expressed in embryonic hypochord where *VEGF* plays a role in the formation of the hypochord (e.g., (Cleaver and Krieg 1998; Cleaver, Seufert, and Krieg 2000; Cleaver et al. 1997)). Our previous work (Senevirathne et al. 2020) showed how the ossifying hypochord may also play a role in modifying the dorsal aorta by occluding it at the posterior-most end of the hypochord and remodeling it to form two branches, which enter the fore- and hind limbs respectively.

### Transcriptomic differences across different time points during urostyle development

The coccygeal and hypochordal tissues chondrify and ossify during development. At the end of metamorphosis, coinciding with the degenerating notochord, they fuse together to form the urostyle. We next delved into identifying genes that switch on/off during metamorphosis and highlight DEGs that are expressed at each time point: before metamorphosis, beginning of metamorphic climax, and end of metamorphosis.

There are numerous studies of metamorphic transcriptomes (e.g., (Brown and Cai 2007; Callery and Elinson 2000; Zhao et al. 2016; Wang et al. 2019; Yaoita and Brown 1990; Kanamori and Brown 1996; Brown et al. 1995)), but none on the urostyle. We first looked into urostyle-responsive transcriptomes by comparing genes that are differentially expressed in the coccyx and hypochord at different time points: 1. Before metamorphosis vs beginning of metamorphosis (electronic supplementary material, figures S1B, S2 and S3) and before/beginning of metamorphosis vs end of metamorphosis (electronic supplementary material, figure S1A). This analysis identified 5664 number of DEGs that fell within the thresholds of FDR< 0.01 (adjusted p-values <0.05 and log fold change of 1.5) and showed unique expression patterns that were significant at each time point.

Several unique sets of genes were up- and down-regulated across the three developmental time points (Fig. S1). Through this step, we identified 4 unique clusters when the transcriptomes were compared between the three developmental time points (before and beginning of metamorphosis *vs* end of metamorphosis (electronic supplementary material, figure S1). Cluster A has 47 genes that were highly downregulated at the end of metamorphosis (“switched off”) compared to the other two time points. This cluster includes genes involved in muscle contraction and M-band stabilization in fast skeletal muscles (e.g., *TRDN* and *MYOM2I*; (Giacomazzi et al. 2017; Auxerre-Plantie et al. 2020)), skeletal development (e.g., *SOX9* (Hattori et al. 2010)), response to inflammation (*PTX3*; (Magrini, Mantovani, and Garlanda 2016)), filament organizing genes (e.g., *KRT18.I* and *VIM.2*; (Velez-delValle et al. 2016; Gan et al. 2016)), extracellular matrix organizing and connective tissue-strengthening (e.g., *COL9A1*, *COL8A1*, *CHAD*; (Brachvogel et al. 2013; Hessle et al. 2014)), and stress regulation (*CRCH.1*; (Reul and Holsboer 2002)). The other two gene clusters, B and C (electronic supplementary material, figure S1A), comprise genes that are both down- and up-regulated at the end of metamorphosis. Cluster C also has 15 genes that are downregulated at the end of metamorphosis, which include collagen markers (e.g., *COL9A3*), and skeletal muscle function genes (e.g., *MYL1* and *ACTN3*; (Schiaffino et al. 2015; Pickering and Kiely 2017)). Genes that are up-regulated (10 genes) are within Cluster B and are involved in mitosis (*CCNB1*; (Strauss et al. 2018)), development of neurons (*POU3F1*; (Zhu et al. 2014)) and maintenance of myelin sheath (*PLP1*; (Gould et al. 2008)). When before metamorphosis was compared with beginning/end of metamorphosis, clustering of the 100 top-most significant genes revealed metamorphic genes that were switched off before metamorphosis but were switched on during metamorphosis. Heatmap clustering revealed five main clusters (electronic supplementary material, figure S1B). Cluster A included 28 genes that were downregulated (switched off) before metamorphosis in both coccyx and hypochord, but as soon as metamorphosis was initiated, these genes were upregulated; they are involved in functions like collagen synthesis (*SERPINH1*; (Widmer et al. 2012)), cell cycle (*CDK6*; (Tigan et al. 2016)), and thyroid hormone inactivation (*DIO3*; (Bianco and da Conceicao 2018)). Cluster B and C includes genes that are switched on prior to metamorphosis and are switched off at the onset of metamorphosis: *HES8*, *FOXP2*, *EGR1*, *HOXD11*, and *PVALB* are representative examples. Cluster D is enrichened with genes that are involved in blood sugar control (e.g., *THRAP3*, *IGF2BP3*; (Choi et al. 2014; Dong et al. 2017)), which are down-regulated before metamorphosis but are up-regulated at the onset of metamorphosis. This part of the transcriptomic analysis identified DEGs that are specific to the three significant time points (before metamorphosis *vs* onset of metamorphosis *vs* end of metamorphic climax). We next explored the GO function of these significant genes during development. The DEGs and the corresponding P-values from the differential expression analyses were imported into an online database of reactome pathways (“Reactome pathway browser”) to compare the functional aspect of these genes (electronic supplementary material, figure S2). DEGs up regulated before metamorphosis were enriched for GO terms like “DNA replication and pre-initiation”, “synthesis of DNA”, “Polymerase switching”, “G1/S transition” (Fig 3.5.B). Whereas the DEGs up regulated during metamorphosis include genes that function in “Collagen formation”, “Cross linking of collagen fibrils”, “*RUNX2* regulated bone development”, and “Osteocyte differentiation” (Fig 3.5.C and D).

Morphological analyses highlighted that both urostyle and hypochord undergo endochondral ossification during development (Senevirathne et al. 2020), and similar ossification patterns were reflected in the gene expression profiles as well. Though there were major differences in some transcriptomes (e.g., presence of T-box genes, *CRCH.1, MMP*s in hypochordal tissues at the onset of metamorphosis *vs* absent in the coccyx), there were similarities in genes that were involved in endochondral ossification: we show that genes that are involved in cartilage and bone formation, extracellular matrix organization, and thyroid hormone responsive elements are present in both tissues (electronic supplementary material, figure S2), but differ temporally (coccyx starts ossifying after 1.5 months, whereas the hypochord initiates its ossification only at the onset of metamorphosis).

### Hypochord, metamorphosis and T-box genes

The ossifying hypochord in anurans is considered an unique feature. As there is no data on the genes that are expressed during hypochordal ossification, we used the DEGs identified by the coccyx *vs* hypochord comparisons (section 3.4.1) to scrutinize this. This analysis identified 470 genes that were uniquely up-regulated only within the hypochordal tissues (they fell within the significant threshold of adjusted p-value <0.05 and FDR<0.01) (Appendix B). Compared to the coccyx, we identified DEGs that were only present in the hypochord (Table 3.2). Out of these, here, we will be focusing on the highly expressed T-box (*TBXT.1*, *TBXT.2*, *TBX1*) genes that are only seen in the hypochordal tissues in this section.

T-box genes have been implicated in early mesodermal patterning and, especially, *Brachyury*/*Xbra* is essential in early mesodermal formation (Cunliffe and Smith 1994, 1992; Hayata et al. 1999; Messenger et al. 2005; Smith et al. 1991), and Brachyury homologues across vertebrates induce the mesoderm (Schulte-Merker and Smith 1995; Yasuoka, Shinzato, and Satoh 2016). *Xenopus* has two paralogues of the gene *Brachyury*: *TBXT.1* (also known at *Xbra* or *T*) and *TBXT.2* (also known as *Xbra3* or *T2*). When *Brachyury* is knocked out, it causes loss of posterior mesoderm and failure to differentiate the notochord (Gentsch et al. 2018; Paraiso et al. 2019). *Brachyury* is also involved in controlling cell fate decisions while acting synergistically with the other transcription factors (like *Bix4*) and genes (*WNT11*) in the posterior mesoderm (Showell, Binder, and Conlon 2004). However, the expression of *TBXT*.1 and *TBXT*.2 in late developing tadpole structures has not been reported so far.

As described below, the temporal and spatial expression patterns of *TBXT.1* and *TBXT.2*, make them good candidate genes for regulating ossification only in hypochordal tissues. To study the potential role of *TBXT.1* and *TBXT.2* in hypochordal ossification further, we performed HCR in-situ hybridization to examine the temporal and spatial expression patterns. *TBXT.1* expression is exclusively concentrated along the ossifying hypochord at the onset of metamorphosis but is not evident in prometamorphic nor at the end of metamorphic climatic tadpoles (Fig. 4).

**Figure 4.**
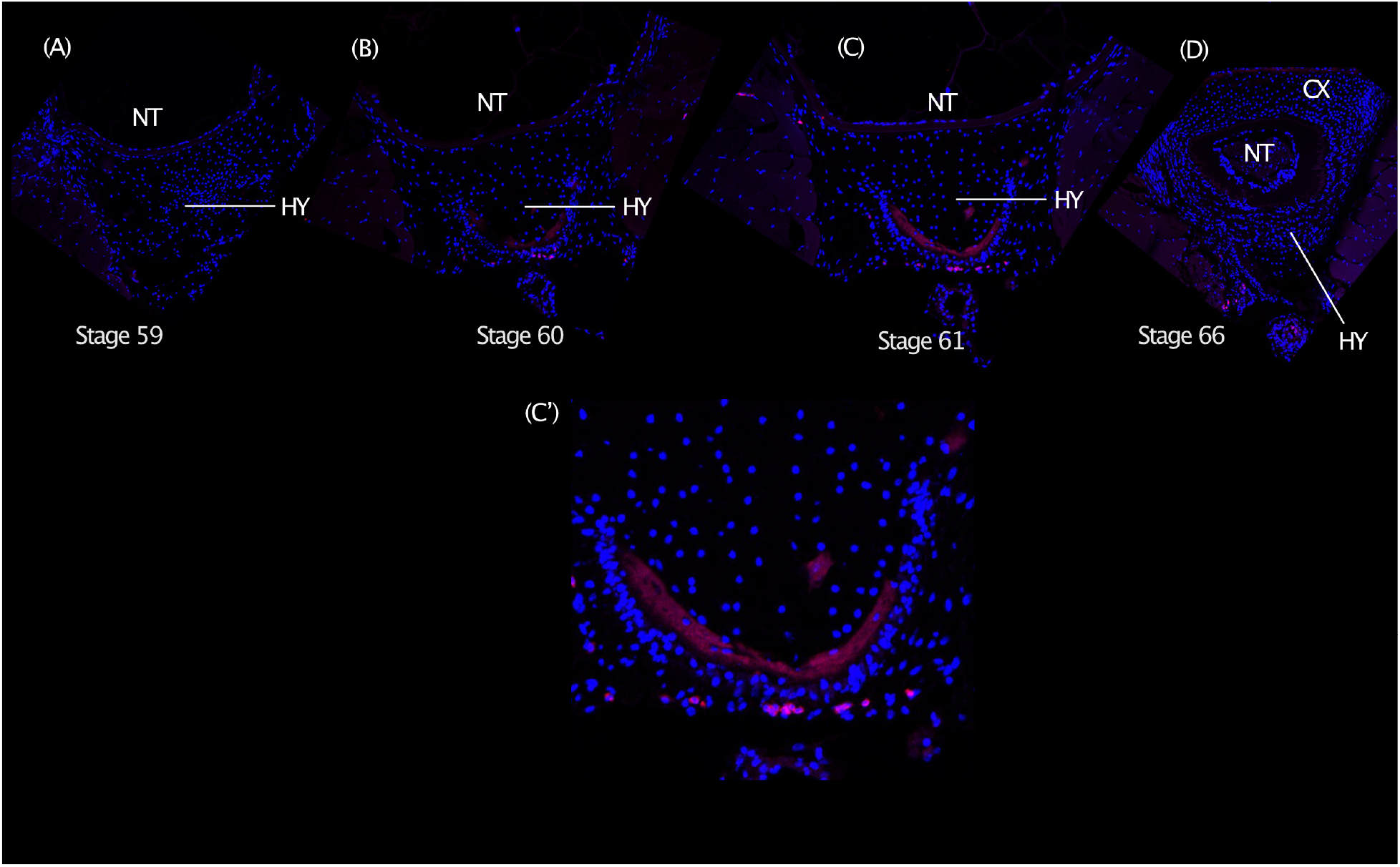
TBXT HCR in-situ hybridization on transverse sections of the urostyle. The periphery of the developing hypochord shows expression of TBXT (pink color), which is initiated once the hypochord starts to form and depletes when the hypochord fuses with the coccyx. Nuclei are stained using DAPI (blue). Abbreviations: CX, coccyx; HY, Hypochord; NT, notochord.

An ossifying hypochord is only normally present in anurans, however, interestingly, hypochord ossification between the caudal part of the vertebral column and notochord also appears as a congenital vertebral anomality seen prenatally in humans, caused by a mutation in the T (*TBXT*) gene (Postma et al. 2014; Ghebranious et al. 2008). In humans with this abnormality, increased expression or duplications of the *TBXT* gene result in production of excess *Brachyury* (Zhu, Kwan, and Mackem 2016; Chen et al. 2020). It has been hypothesized that this excess *Brachyury* causes residual cells ventral to the notochord to grow and ossify in humans and sometimes results in sacral agenesis in newly born babies (commonly to referred to as the “frog-like” syndrome). The observation of high levels of *TBXT*/*TBXT2* in ossifying hypochordal cells (which is ossified ventral to the notochord) and presence of two duplicated copies of the *TBXT* (*T* and *T2*/*TBXT* and *TBXT.2*) in anurans compared to normal humans and other vertebrates is thus tantalizing and needs further scrutiny. Previous studies have shown that *Brachyury* acts as a switch in posterior mesoderm specification during embryogenesis and is restricted to the anteroposterior axis (Cunliffe and Smith 1994, 1992). Here, during hypochordal ossification, the onset of metamorphosis could be triggering ectopic expression of *TBXT*/*TBXT.2* in hypochordal cells, which could potentially express posterior mesodermal genes and subsequently activate down-stream targets of *TBXT*/*TBXT2*, which in turn initiates chondrification and ossification.

### Transcriptomic comparisons between coccyx + hypochord and other ossifying elements

Vertebrate ossification happens by two major processes: endochondral (cartilaginous precursors used as a template) and intramembranous (direct ossification of the condensed mesenchymal cells) (Breeland, Sinkler, and Menezes 2021). Even though coccyx and hypochord are derived from two different cell populations, they both undergo endochondral ossification (Senevirathne et al. 2020). During this process, mesenchymal cells condense (commit to form osteoprogenitors) and aggregate to form cartilaginous precursors during early development. Cartilaginous precursors expand and cells proliferate, next the extracellular matrix is synthesized, and finally, mineralization of the matrix occurs. These steps are similar to other bones in vertebrates, which undergo endochondral ossification as well (Mackie et al. 2008). However, to see if the transcriptomic profile during this process is conserved in the two bones that form the urostyle, we compared the spatial and temporal transcriptomic maps of the osteocytes (from published datasets of different skeletal tissues of different ages) with my current dataset.

Youlten et al. (Youlten et al. 2021) identified three clusters of gene ontology (GO) functions during osteocyte development: 1. “An early expression cluster” (expressed in osteoprogenitors/osteoblast-like cells); 2. “An early activation cluster” (expressed in early osteocytes); 3. “A maturation cluster” (expressed in mature osteocytes). We compared the expression of the genes belonging to these GO functions with the coccygeal and hypochordal transcriptomics to see if the molecular underpinning of ossification is similar in the genes responsible for the formation of the urostyle as well.

#### Early expression cluster

This included GO term functions “Extracellular matrix organization”, “Angiogenesis”, “Cartilage development”, and “Connective tissue development” (electronic supplementary material, figure S4). Out of the genes that are differentially expressed, there are some that are inactive before metamorphosis in the hypochord (e.g., *COL22A1*, *COL16A1*, *COL6A3*, *RUNX1*, *IHH*), but are highly expressed once the metamorphosis is initiated. High expression of these genes in the coccygeal cells even before the onset of metamorphosis corroborates our morphological studies, where we revealed that the post caudal vertebrae of the coccyx initiated mesenchymal cell aggregation early in development (1.5 months after embryogenesis) *vs* 2 months in hypochord. Apart from the differences in the temporal expression of genes within the “Early expression cluster”, a few genes involved in cartilage development are not present in the hypochord compared to the coccyx (e.g, *FOXL1*, *RUNX3*, *FOXD3*, *PMM2*, *EDN1*).

#### Early activation cluster

This cluster includes the GO terms “Axon guidance”, “Axon development”, “Axogenesis”, “Regulation of axogenesis”, and “Neuron projection guidance” (Fig. 5). While the coccyx DEGs act in a similar way to the rest of the long bones in vertebrates within this cluster, hypochord shows a different pattern. Most of the genes (e.g., *NTRN*, *SLITRK3*, *POUF42*, *DCC*) that are discussed as essential regulators in guiding the axons in long bones are not expressed within the hypochord (Fig. 5).

**Figure 5:**
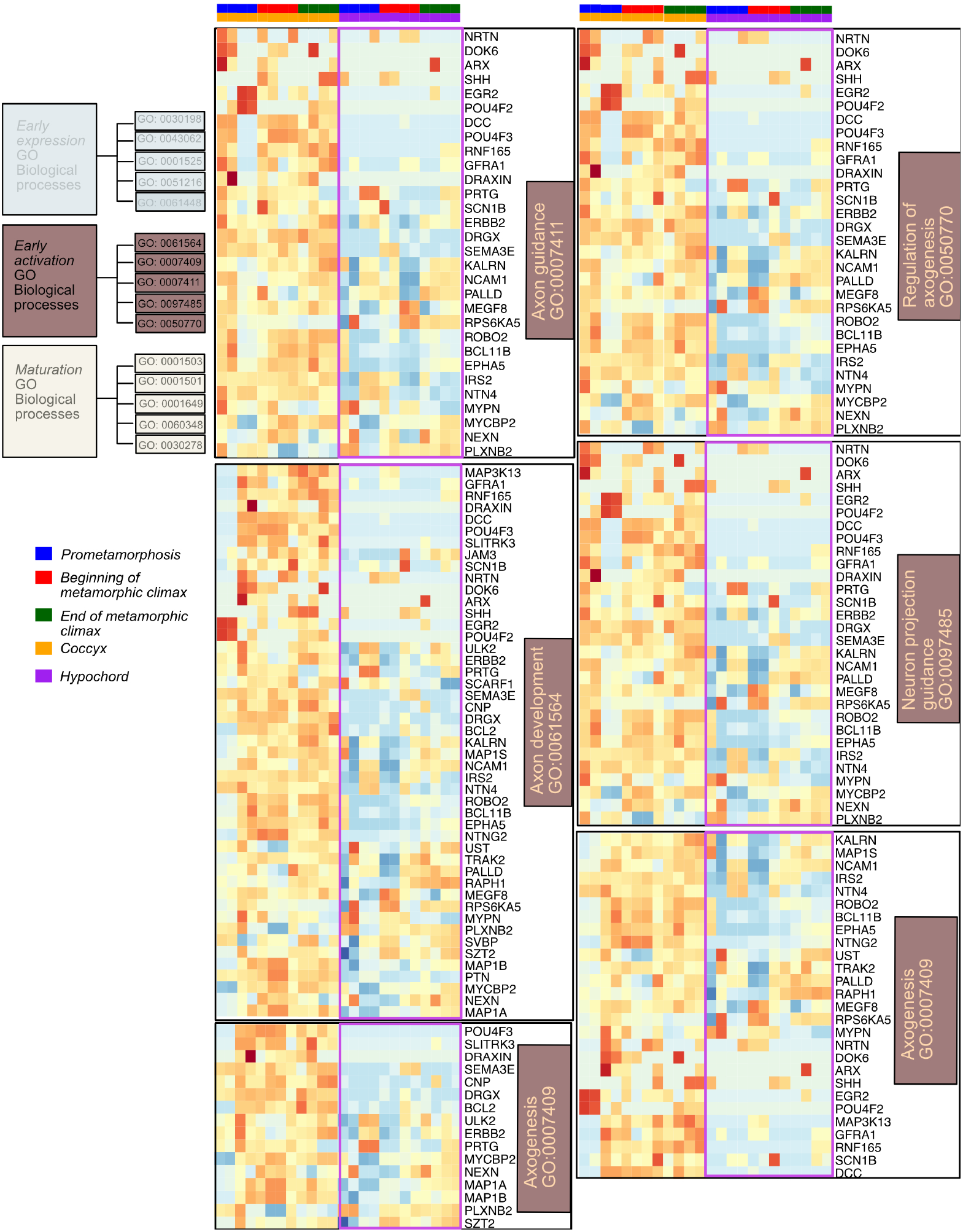
Heatmaps showing differentially expressed genes involved in GO functions belonging to the “Early activation cluster” of osteocyte differentiation. Significant genes of the osteocyte transcriptome are divided into three clusters *(Youlten et al. 2021)*. This cluster includes the GO functions Axon guidance, Axon development, Axogenesis, Regulation of axogenesis, and Neuron projection guidance. Genes of interest that are differentially expressed between the coccyx and hypochord are highlighted in purple color.

#### Maturation cluster

The GO term functions “Bone development”, “Skeletal system development”, “Regulation of ossification”, “Ossification”, “Osteoblast differentiation” are included in this cluster (supplementary material, figure S5). Maturation period in the hypochord happens once the metamorphosis is initiated and when the tadpole reaches the end of its metamorphic climax (supplementary material, figure S5). Within the hypochord, genes involved in ossification (e.g., *GPC3*, *TMEM19*, *IFITM5*, *COL11A1*, *PHOSPHO1*, *SOX8*) and osteoblast differentiation (e.g., *GLI1*, *FBN2*, *SATB2*) are highly expressed in tadpoles at the end of the metamorphic climatic and are inactive at prometamorphic stages. Comparatively, in the coccyx, since the ossification happens prior to the metamorphic climax, the majority of the genes are highly expressed even at the beginning of metamorphosis. A few genes (e.g., *TBX15*, *BARX2*, *SHH*, *AXIN2*) are not expressed in hypochord nor in the coccyx, compared to the other ossifying long bones in vertebrates.

This transcriptomic comparison led to three main findings: (1). Between the two tissue types, the coccyx’s DEGs share similarities with the other bones’ transcriptomics in vertebrates. (2). Hypochord undergoes its early activation period before metamorphosis, and a maturation period once metamorphosis is initiated. (3) Hypochordal DEGs lack an early activation period, which includes most of the axon developing genes.

### ATAC-seq and Urostyle-responsive gene regulation

During anuran metamorphosis, the larval body form undergoes dramatic remodeling within 6-8 days, and this is reflected in both morphological and gene expression patterns. Therefore, it can be extrapolated that gene regulation changes over this same time period. To study the underlying changes in chromatin accessibility, we used an ATAC-seq approach using the same developmental stages and the same number of replicates as the RNA-seq work. The number of peaks varied between the three stages that we used: before metamorphosis (4563 peaks), beginning of metamorphosis (6805 peaks), and end of metamorphosis (6805 peaks). More than 50% of peaks were distributed in distal intergenic regions. The rest of the peaks were distributed along intronic, exons, and promoter regions. When comparing the three time points, the most significant change of peak distribution observed was the percentage of peaks that fell on the exon regions (other than the 1^st^ exon): before metamorphosis the percentage was lower (<1%) when compared with the number of peaks that were seen at the beginning and at the end of metamorphosis (7–10%) (Fig. 6B).

**Figure 6:**
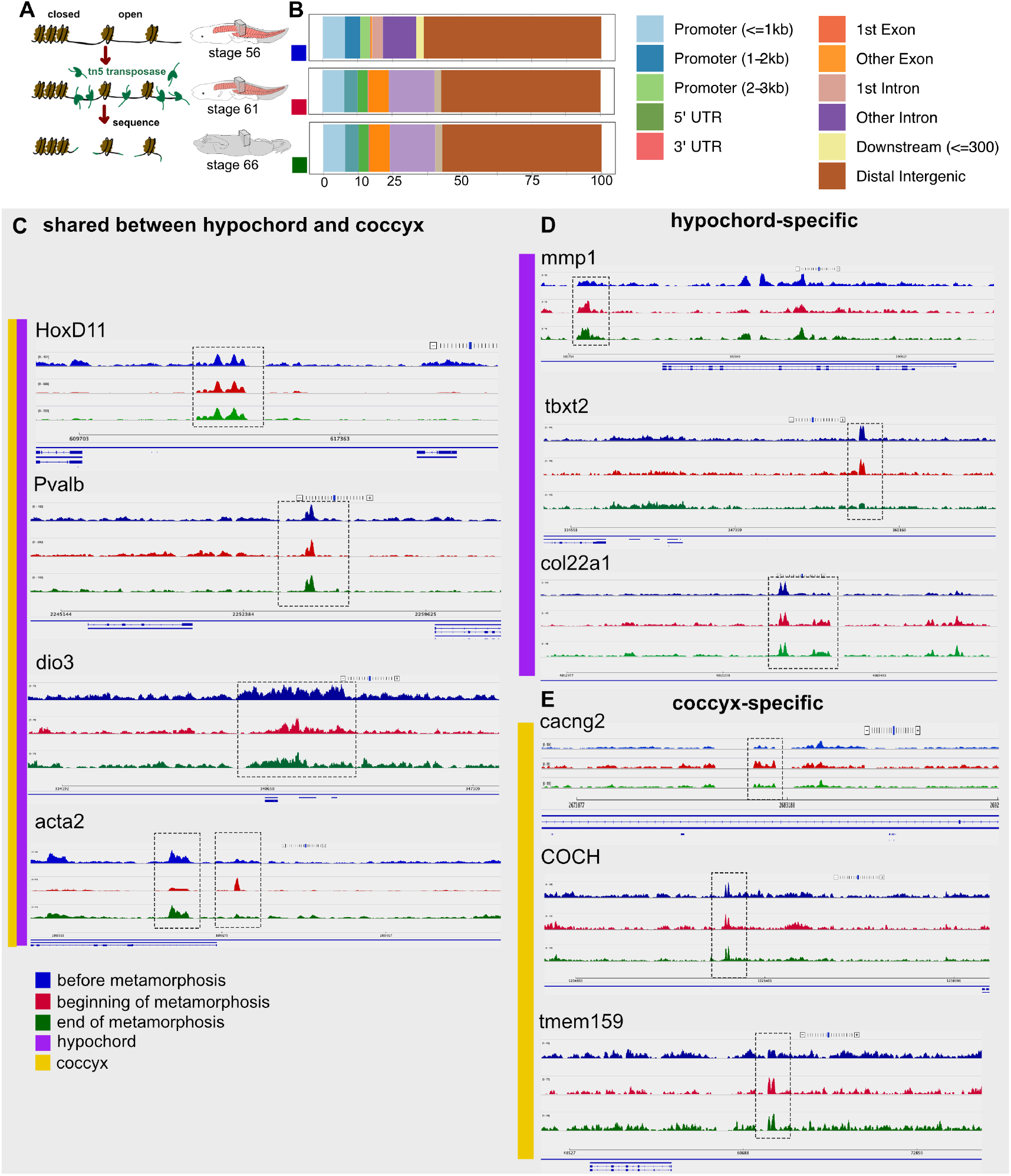
Urostyle-responsive regulatory regions. A. Schematic diagram showing the workflow for chromatin profiling experiment. B. Proportions of developing urostyle ATAC-seq peaks annotated to different genomic regions across development; majority of the peaks fall within the distal intergenic region and beginning (stage 61) and end of metamorphic climatic (stage 65) peaks differ from the prometamorphic (stage 56) ATAC-seq peaks with respect to peaks falling within the exon regions that are not the first exon. C–E. ATAC-seq urostyle profiles at stage 56 (blue), stage 61 (red), and stage 65 (green) at the loci of validated up-regulatory genes narrowed down from RNA-seq analyses.

Next, we compared the ATAC-seq data with the RNA-seq data and observed that majority of the peaks are located close to the up-regulated genes in the hypochord and coccyx that were identified from the transcriptomic data. The genes *TBXT* and *TBXT.2*, which are up regulated in the hypochord, have peaks located within the intronic regions before and at the beginning of metamorphosis, and the peak is lost at the end of metamorphosis (Fig. 6D). Other genes expressed in hypochordal tissues like *MMP1* and *COL22A1* have peaks downstream of the genes and are seen only once the metamorphosis is initiated. Genes that were upregulated in the coccyx, e.g., *HOXD11*, *PVALB*, *DIO3*, and *ACTA2* have ATAC-seq peaks closer to each gene and were present throughout development (Fig. 6C–E). This could be because the coccygeal ossification occurs early in development (after 1.5 months) compared to the hypochord. These results highlight urostyle-responsive regulatory regions during development and need further scrutinization using functional assays.

## DISCUSSION

The anuran urostyle, composed of a coccyx and a hypochord, reflects how novel structures facilitate evolution of new body plans. Our previous work presented a morphological analysis of the ontogeny of the anuran urostyle using immunohistochemistry, histology, bone and cartilage staining, and microCT scanning. Through this, we identified cells of interest and the developmental stages to target for this follow up study. To elucidate how this structural novelty arose and its genetic underpinnings, we used a spatial transcriptomic (RNA-seq) and an ATAC-seq approach.

### T-box genes and the hypochord

The coccyx and hypochord have two sets of differentially expressed genes. Hypochordal genes are active at the onset of metamorphosis, whereas the coccygeal DEGs are highly expressed even before metamorphosis. This analysis revealed a large set of genes (Tables 3.1 and 3.2 and Appendices C, D) that are uniquely up regulated in the hypochord and have not been reported before. One of the most significant groups of genes that is upregulated in the hypochord are the T-box genes (*TBXT* and *TBXT.2*). T-box genes have a 180-bp DNA binding domain that is highly conserved. Orthologues of the gene *Brachyury*, one of the highly expressed T-box genes in the hypochord, are present in all multicellular organisms (Chen et al. 2020). *Brachyury* is important in posterior mesoderm development (initially expressed in the developing mesoderm but later restricted to the tail bud and notochord) (Hotta et al. 2000). While early mesoderm differentiation patterning depends highly on *TBXT*/*TBXT.2*, a role for these genes in later developmental stages has not been previously reported or discussed. During metamorphosis, the tadpole body undergoes dramatic remodeling, including tail loss and development of new structures like the urostyle. The hypochord, thought to be of an endodermal or superficial mesoderm origin, undergoes ossification at the beginning of metamorphosis only in anurans. We hypothesize that presence of high levels of *TBXT*/*TBXT.2* causes the hypochordal cells to undergo ossification at the onset of metamorphosis. Such unusual ossification appears to also occur in response to a congenital vertebral column malformation (VCM) in humans that happens because of a *Brachyury* gene mutation in the intron 7 (Ghebranious et al. 2008) and in the highly conserved T-box sequence (Postma et al. 2014); these VCMs eventually lead to sacral agenesis (“frog-like”) syndrome in babies. Apart from these mutations, *TBXT*/*TBXT.2* genes also induce EMT in humans when over expressed in carcinoma cells (Henderson et al. 2005), and it has also been recorded that duplications of the *Brachyury* gene cause vertebral column chordomas (Vujovic et al. 2006; Henderson et al. 2005). Frogs have two paralogues of *Brachyury* genes, perhaps explaining the overexpression of *TBXT*/*TBXT.2* at the onset of metamorphosis, which could in turn allow the T genes to activate downstream targets that lead to chondrification and ossification. When *Brachyury* genes are highly expressed in human chordoma cells, matrix metalloproteinases (e.g., *MMP12*, *MMP13*, *MMP24*) (Wan et al. 2016) are also upregulated at the same time (which is also seen in hypochordal cells). The extent to which the human and frog conditions are similar awaits functional tests.

### Coccyx and hypochord vs other vertebrate skeletal elements

Coccyx and hypochord undergo endochondral ossification and show an array of genes that are similar to the genes expressed in other long bones that undergo endochondral ossification in vertebrates (e.g., mesenchymal-to-chondrocytes involved genes like *BMP*s, *SOX9*; chondrocytes-to-osteoblasts/osteocytes was seen in highly expressed genes like *RUNX2*, Osterix, *IHH*). Apart from these similarities, when comparing the already published osteocyte transcriptomics (Youlten et al. 2021), hypochord shows some considerable differences among the rest of the bones in vertebrates. Hypochordal cells express osteoprogenitor-specific genes before the metamorphic climax, and metamorphosis acts as a switch that activates osteogenesis (*vs* in coccyx osteogenesis is initiated prior to metamorphosis). Other than the temporal differences observed regarding ossification, the DEGs of the hypochord reveal that hypochordal cells lack the “early activation phase,” which includes regulators needed in “Axogenesis” and “Axon development” in ossifying bones (Fig. 5). Vertebrate bones are innervated by sensory and sympathetic nerves during skeleton development (Tomlinson et al. 2020), where the periosteum and bone marrow have the highest density of nerves whereas the mineralized matrix has very few (Mach et al. 2002; Castaneda-Corral et al. 2011; Tomlinson et al. 2020). During development, bone innervation and endochondral ossification happen simultaneously (Tomlinson et al. 2020), and it is hypothesized that axon guidance regulates formation of the neuronal network, which is subsequently required for the osteocyte network formation (Youlten et al. 2021). It is surprising that the ossifying hypochord lacks the genes needed for axon development (Fig. 5), and our results raises the possibility that the hypochordal development maybe disconnected from the neuronal signals. Future work is needed scrutinizing the innervation patterns within the hypochord during its development to better understand this.

Our integrative approach, using morphological and molecular data sets (genes and gene regulation) on the development of the urostyle, scrutinizes the evolution of a novelty. This has been evolutionary favored for more than 200 million years and is seen in all extant anurans during their development. We propose that the underlying changes in the genetic network gave rise to the anuran urostyle, and it is an evolutionary novelty that has enabled successful inhabitation of several ecological niches. Future work targeting the candidate genes responsible for the development of the urostyle, together with functional assays, will shed light on the evolution of this structural enigma.

## ACKNOWLEDGMENTS

We would like to thank Marko Horb, Nikko-Ideen Shaidaini, and Marcin Wlizla (National Xenopus Resource [NXR], Marine Biological Laboratory) for husbandry and providing *X. tropicalis* tadpoles; James Hanken, Victoria Prince and Shubin Lab members for their comments and helpful discussions on this work. This work was supported by University of Chicago Biological Sciences and the Brinson Foundation (to N.H.S.) and by O’Brien and Hasten Fellowship to G.S.

## AUTHOR CONTRIBUTIONS

G.S. and N.H.S conceptualized the project and designed research. G.S. Performed research and analyzed data. G.S. wrote the paper with inputs from N.H.S.

## Supplementary Material

**Figure S1:**
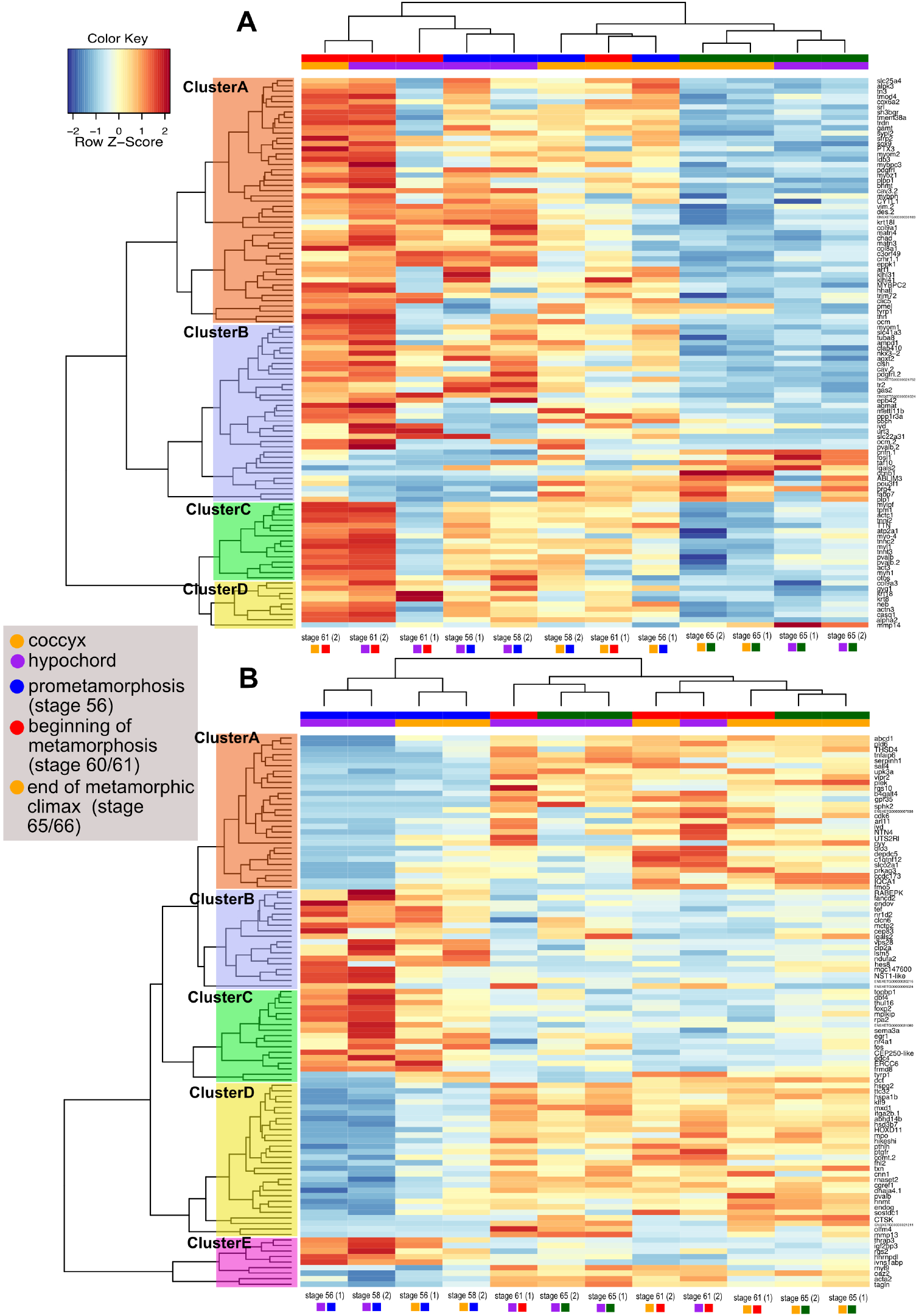
Differentially expressed genes across different time points during urostyle development. A. Heatmap showing before and beginning of metamorphosis vs end of metamorphosis, indicating that there is a set of genes (eg., *PTX3*, *SOX9*, *KRT18*) that switch off at the end of metamorphosis in both tissue types. B. Heatmap comparing before and beginning of metamorphosis, highlighting a set of genes (eg., *DIO3*, *HOXD11*, *PVALB*) that switch on during urostyle development. Purple: coccyx; Yellow: hypochord; Blue: before metamorphosis; Red: beginning of metamorphosis; Green: end of Metamorphosis

**Figure S2:**
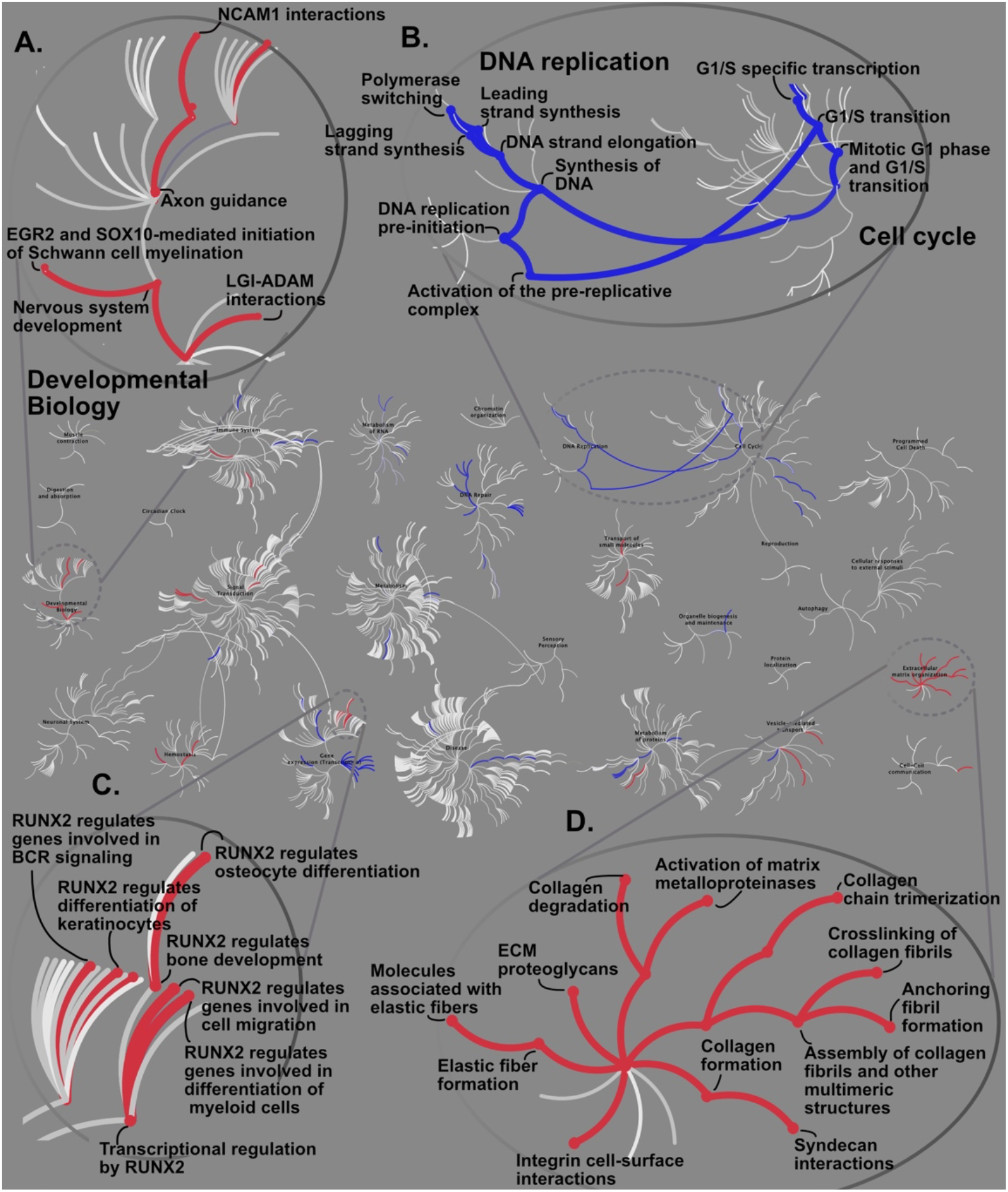
Reactome pathway analysis for up/down regulatory genes in two different time points: before metamorphosis (blue) and beginning of metamorphosis (red). The central circles represent a top-level pathway, and the circles away from the center represents lower levels in each respective pathway. Zoomed-in sections of top-level pathways of Developmental Biology (A), DNA replication and Cell cycle (B), Gene expression (C), and Extracellular matrix organization (D) are shown. Overrepresented pathways (P < 0.05) are colored in blue (prometamorphosis) and red (beginning of metamorphosis). Pathways that are not significant are shown in light gray lines.

**Figure S3:**
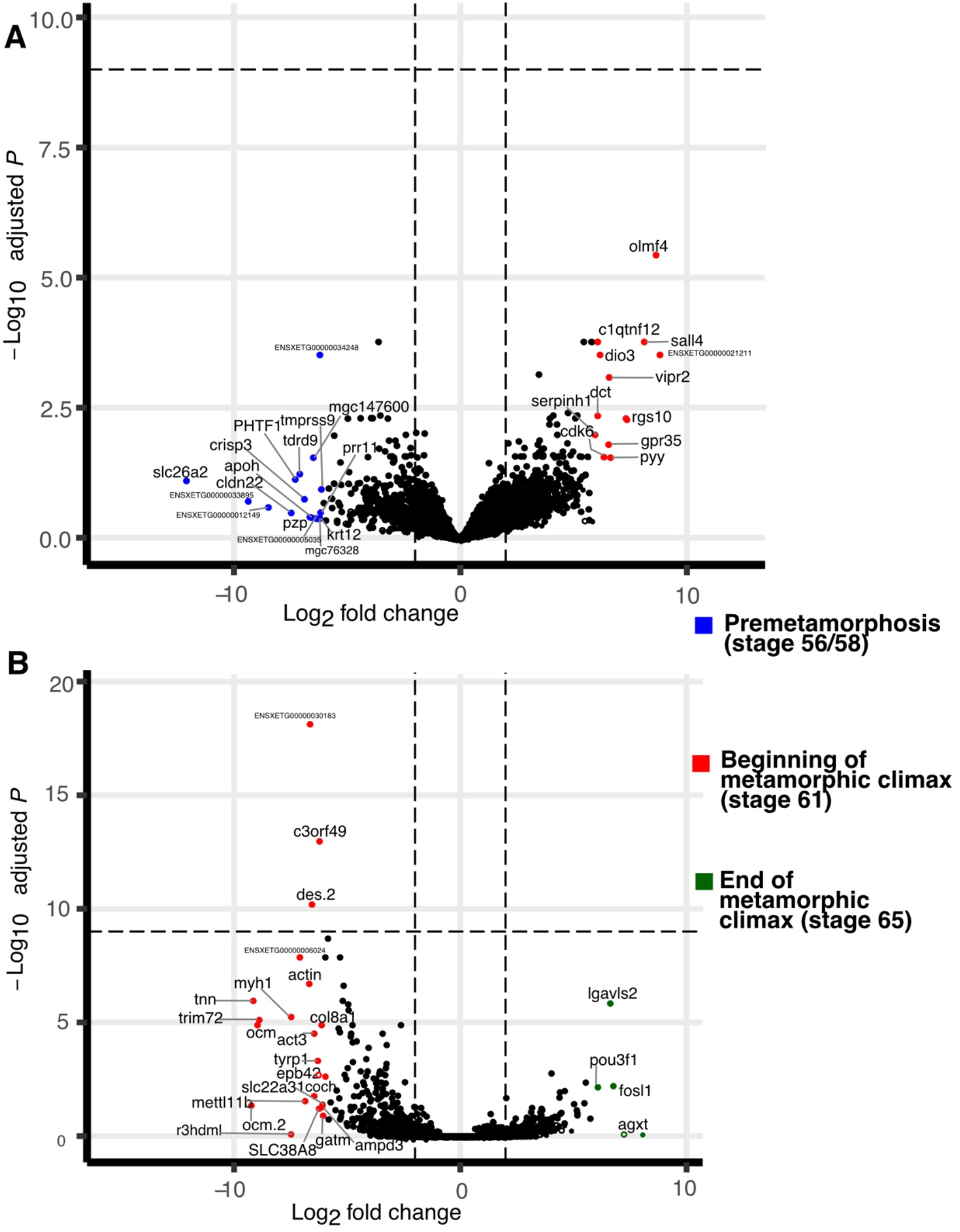
Volcano plot showing differentially expressed genes across three developmental time points, during the formation of the urostyle (P < 0.05, FDR<0.01).

**Figure S4:**
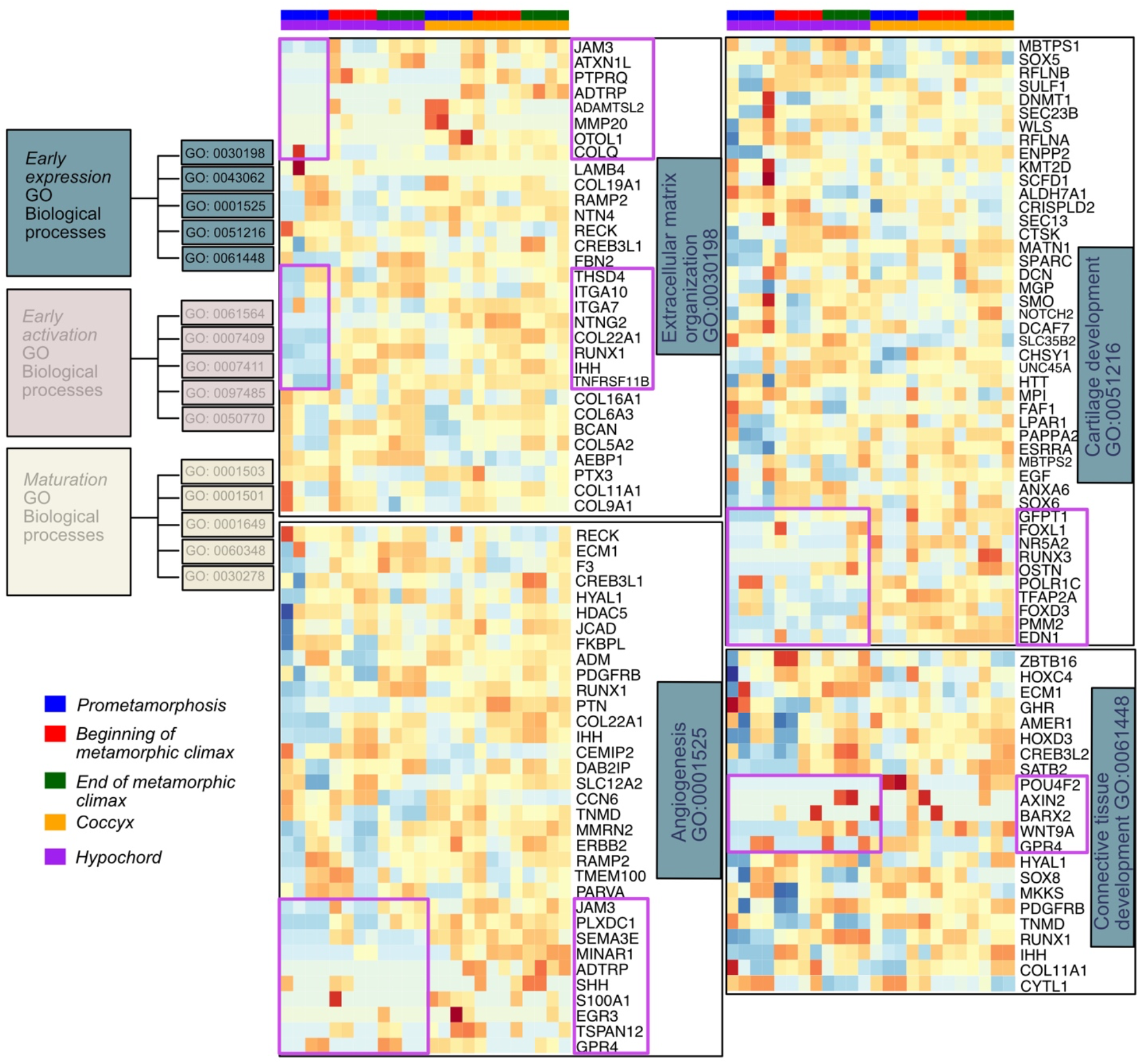
Heatmaps showing differentially expressed genes involved in GO functions belonging to the “Early expression cluster” of osteocyte differentiation. Significant genes of the osteocyte transcriptome are divided into three clusters (Youlten et al. 2021). This cluster includes the GO functions Extracellular matrix organization, Angiogenesis, Cartilage development, and Connective tissue development. Genes of interest that are differentially expressed between the coccyx and hypochord are highlighted in purple color.

**Figure S5:**
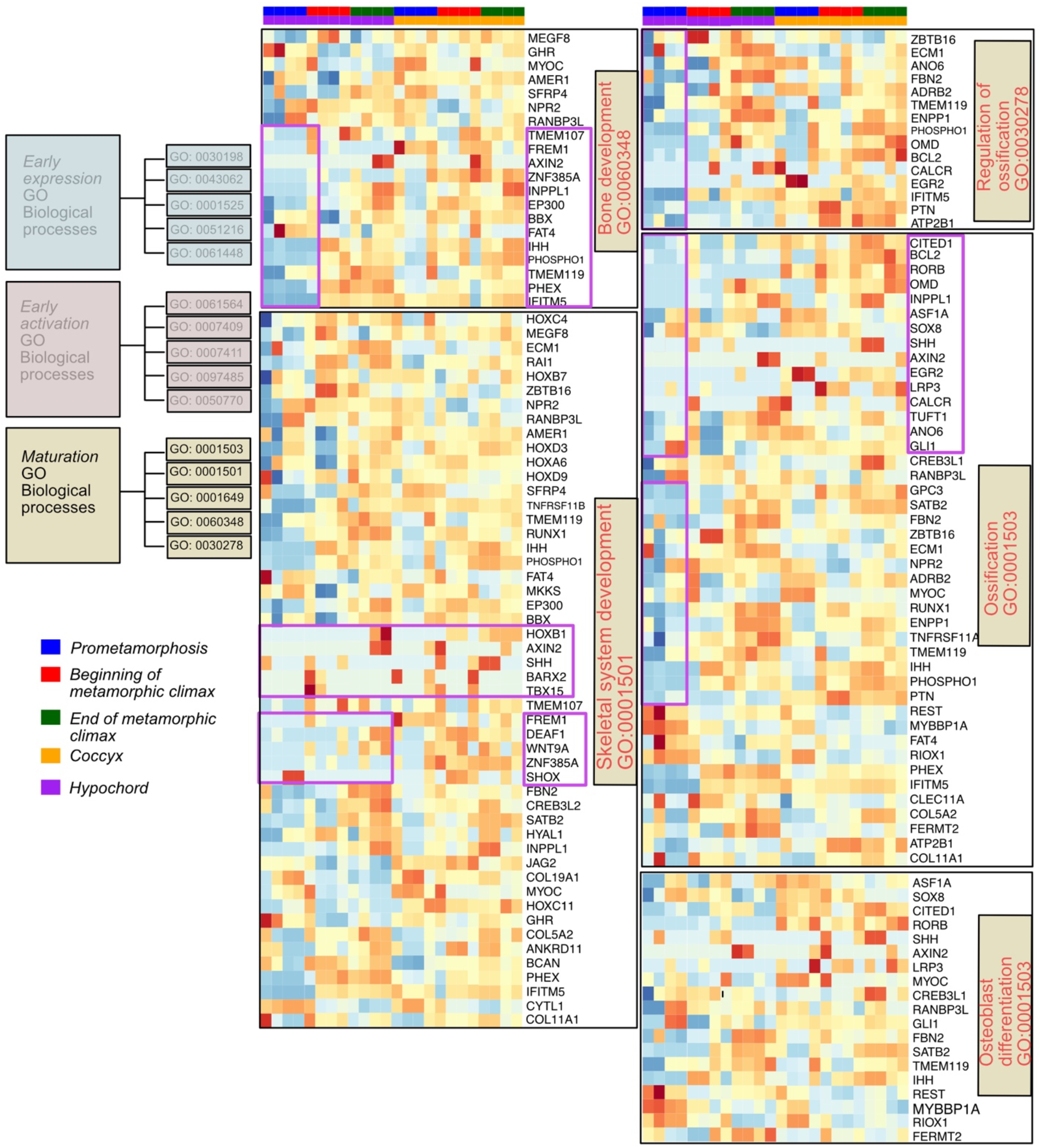
Heatmaps showing differentially expressed genes involved in GO functions belonging to the “Maturation cluster” of osteocyte differentiation. Significant genes of the osteocyte transcriptome are divided into three clusters (Youlten et al. 2021). This cluster includes the GO functions Bone development, Skeletal system development, Osteoblast differentiation, Ossification, and Regulation of ossification. Genes of interest that are differentially expressed between the coccyx and hypochord are highlighted in purple color.

